# Distinct mediodorsal-prefrontal loops differentially encode reward-predictive cues

**DOI:** 10.64898/2026.03.02.709125

**Authors:** Kelly Runyon, Kyra Sanders, Alec Hartle, William Matt Howe

**Affiliations:** School of Neuroscience, Virginia Tech, Blacksburg, Virginia, United States

## Abstract

Using external cues to guide behavior is a core function that enables multiple aspects of cognition and attentional control, and deficits in this process are central to many theories of neuro-psychiatric, degenerative, and developmental disorders. Cue detection relies on the precise coordination of neural circuits, with the mediodorsal thalamus (MD) hypothesized to play a pivotal role in orchestrating the relay of cue-based associative information to the prefrontal cortex. The prefrontal cortex comprises multiple subregions, which are believed to differentially contribute to such associative cue-based behaviors. This regional specificity is likely seeded by projection-defined MD→PFC pathways, although the anatomical organization of these discrete channels and their dynamic roles in cue detection are still being defined. Here, we address this gap by combining anatomical circuit mapping of MD-PFC output pathways with in vivo calcium imaging during a cue-based reward conditioning task in mice. These experiments reveal that MD projections to distinct PFC subregions (prelimbic and anterior cingulate cortex) form topographically defined loops, that are characterized by unique patterns of activity across cue-reward learning. Using fiber photometry to monitor changes in calcium activity in axonal projections from the MD to the PFC, we show that during learning, MD projections to the prelimbic subregion are activated by cue presentation, and the dynamics of this activity remain stable across training days. In contrast, MD projections to the anterior cingulate exhibit a learning-dependent suppression of activity that predicts reward approach behavior in late training. Interestingly, the two pathways exhibit opposing activity patterns when the predictive validity of the cue is diminished by extinction training, suggesting distinct functional roles in detecting violations of learned contingencies. Together, these findings reveal previously unrecognized anatomical and functional distinctions within MD-PFC circuits and demonstrate that parallel thalamocortical pathways differentially support cue detection and behavioral flexibility. This work advances understanding of thalamocortical mechanisms underlying cue detection and may inform circuit-based approaches for treating cognitive dysfunction in psychiatric disorders.

## Summary

The capacity to detect predictive cues and use them to guide future behavior is a fundamental component of decision making, and disruption of this process is a hallmark of many neurodevelopmental and neuropsychiatric disorders (Ouhaz et al., 2018; Welsh et al., 2010). mediodorsal thalamus (MD) is a higher-order thalamic nucleus that shares reciprocal connections with the prefrontal cortex (PFC), and these circuits are essential for such cue-guided behavior (Goldman-Rakic & Porrino, 1985; Hasselmo & Sarter, 2011; Miller & Cohen, 2001). Across all species, the prefrontal cortex exhibits functional topography, and in the rodent subregions like the prelimbic (PRL) cortex and anterior cingulate cortex (ACC) are believed to differentially contribute to use of learned cue-outcome relationships (Anderson et al., 2011; Balleine & Dickinson, 1998; Gritton et al., 2016; Howe et al., 2017; Sharpe & Killcross, 2015) and allocation of attentional resources based on expectations and feedback (Dalley et al., 2004; Devinsky et al., 1995), respectively. While the MD projects to each of these regions, the specific organization of MD→PRL and MD→ACC pathways, and their contributions to the functional topography of the PFC, are not clear. Although MD–PFC connectivity has been reviewed extensively (Ferguson & Delevich, 2020; Parnaudeau et al., 2018; Wolff & Halassa, 2024), the anatomical segregation and functional roles of MD→PRL and MD→ACC pathways have not been systematically examined. To better understand how these pathways may support different components of cue-guided behavior mediated by the PRL and ACC, here we combined retrograde neuronal tracing and calcium imaging of MD axonal inputs to these discrete PFC subregions in mice undergoing cue-reward conditioning.

## Results

Although classical views have focused on the medial, central, and lateral subdivisions of the MD (Sarter & Markowitsch, 1984), recent findings suggest that thalamic nuclei exhibit anatomical and functional specificity along the anterior-posterior axis (Barreiros et al., 2021; Mandelbaum et al., 2019). Therefore, we first used the retrograde tracer CTb to explore the organization of MD projections to the PRL and ACC across each of these levels of organization (Figure 1A). In addition, because the MD forms reciprocal loops with the PFC (Parnaudeau et al., 2018; Xiao et al., 2009) we assessed the organization of PRL and ACC projection to the MD (Figure 1B). Finally, to assess the functional dynamics of these circuits during behavior, we expressed GCaMP7s in MD and measured calcium activity in MD axon terminals in PRL or ACC during a cue detection task (Figure 1C-H).

**Figure 1.**
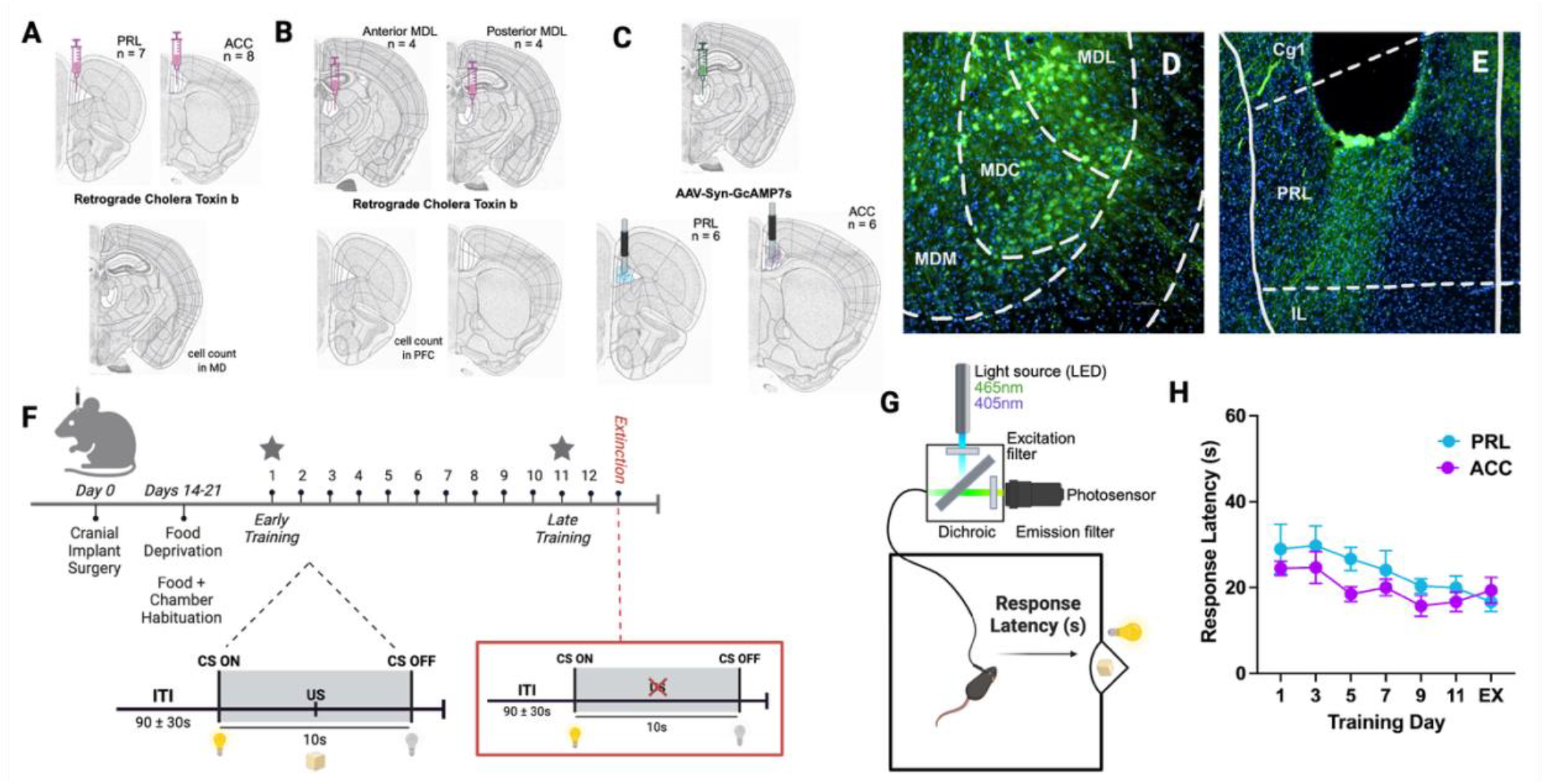
Overview of experimental design: retrograde tracing, MD axon terminal recordings, and cue detection task used to characterize MD→PRL and MD→ACC circuit anatomy and dynamics. (A) To identify mediodorsal thalamus (MD) neuron populations projecting to prelimbic cortex (PRL) or anterior cingulate cortex (ACC), the retrograde tracer cholera toxin subunit B (CTb) was injected unilaterally into PRL (n = 7) or ACC (n = 8) of C57BL/6 mice. Labeled neurons were quantified across the anterior-posterior axis of MD. (B) To assess reciprocal connectivity, CTb was injected into either anterior or posterior MD (anterior group, n = 4; posterior group, n = 4), and retrogradely labeled neurons were quantified in PRL and ACC, allowing examination of PFC inputs to MD. (C) To explore the function of these two MD projections (PRL, ACC) during cue detection and utilization, GCaMP7s was expressed in MD and a unilateral optical fiber was implanted over PRL or ACC to measure calcium activity in MD axon terminals projecting to each region (PRL n = 6; ACC n = 6). (D) Representative image showing GCaMP7s virus expression in MD. (E) Representative image showing optical fiber placement above PRL. (F) Schematic of the cue detection task, in which a 10-s light cue (CS) predicts sucrose pellet delivery at 5s (US). (G) Experimental timeline showing 12 days of training followed by one day of cue-reward extinction, during which projection-specific calcium signals were recorded using fiber photometry. First and last days of recording were used as timepoints of interest. (H) Behavioral performance measured as response latency-the time from cue onset to reward port entry-reflecting learned cue-reward associations (Robinson & Flagel, 2009). Latency remained stable across training and extinction days.

Beginning with the anatomical organization of MD projections to prefrontal cortex, we first quantified retrogradely labeled neurons in the MD following CTb injections into either PRL or ACC (Figure 2A-C; Nissl atlas images from the Allen Reference Atlas – Mouse Brain (Science, 2011). Consistent with prior reports (Wolff & Halassa, 2024), CTb injections in PRL and ACC yielded retrogradely labeled neurons that were largely confined to the MDL. This initial quantification was based on fluorescence intensity within each MD subregion, normalized to background signal, rather than individual cell counts. A linear mixed-effects model revealed a significant main effect of MD subregion (*F*(2, 52) = 3.59, *p* = 0.035), reflecting greater labeling in lateral relative to central and medial MD. Having established that projections were concentrated within the MDL, we next examined whether PRL- and ACC-projecting neurons were differentially organized along the anterior-posterior axis within this subregion. Models revealed a significant interaction between PFC region (PRL vs ACC) and labeled neuron density along the anterior-posterior axis in MDL. Specifically, MD neurons projecting to PRL were concentrated in the anterior portion of the MDL, whereas neurons projecting to ACC were enriched in the posterior MDL (Figure 2A-C). Analyses of labeled neurons in the PRL and ACC following CTb infusions into either anterior MDL or posterior MDL indicated a significant interaction between injection site and PFC subregion, such that PRL predominantly targeted the anterior, and ACC posterior, MDL (Figure 2D-E). Together, these results are in line with previous descriptions of MD-PFC connectivity (Collins et al., 2018; Sherman, 2017), here shown to extend to here shown to extend to a topographically organized, reciprocal loop architecture between anterior MDL-PRL and posterior MDL-ACC.

**Figure 2.**
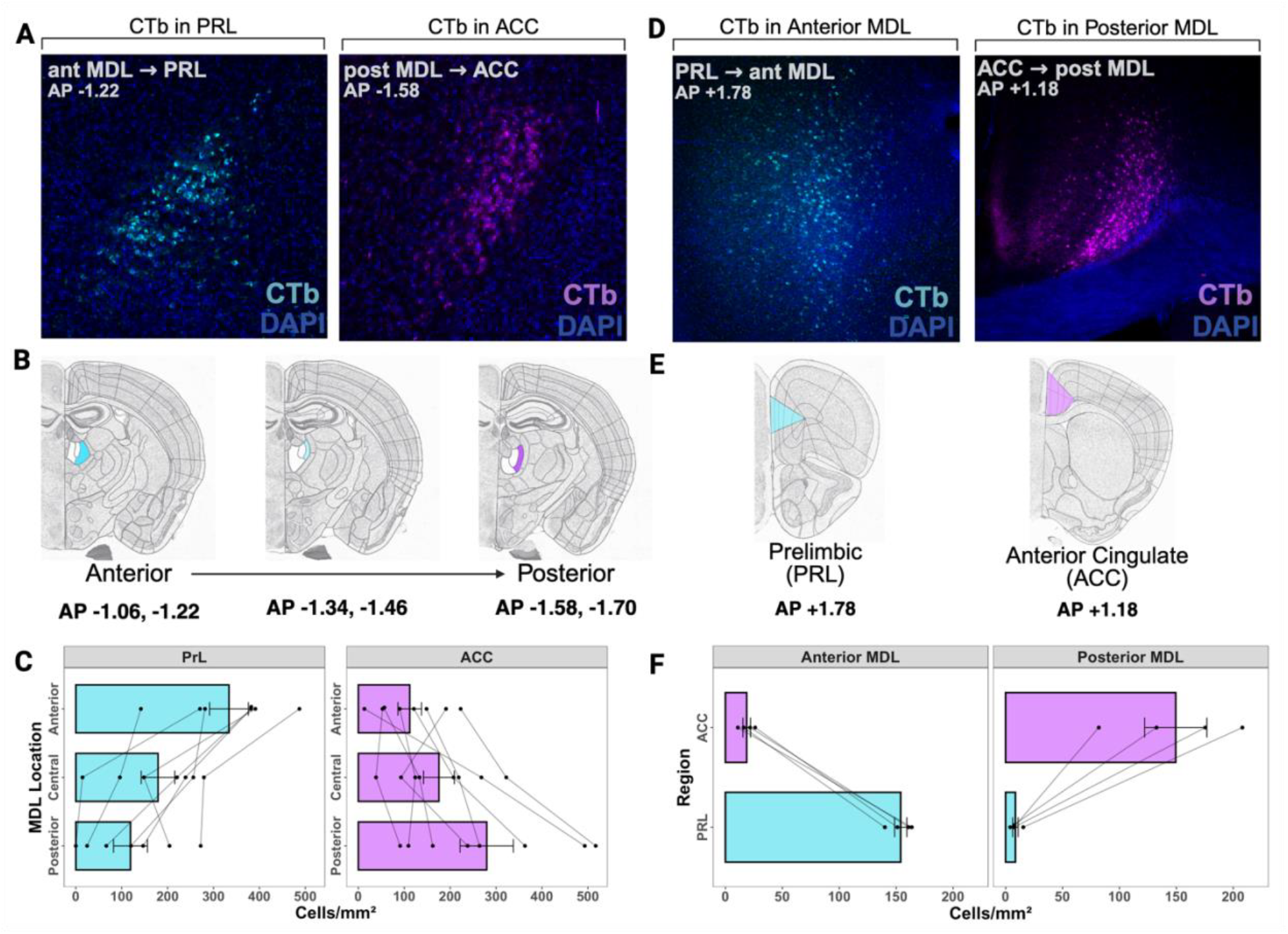
MD ↔ PFC projection anatomy reveals distinct anterior-posterior organization and reciprocal structural loops. (A) Retrograde tracer (CTb) injections into PRL (n = 7) or ACC (n = 8). Representative tissue images show labeled neurons in (left to right) anterior MDL and posterior MDL. (B) Nissl atlas images from the Allen Reference Atlas - Mouse Brain showing anterior-posterior groups. (C) LME with injection site as a between-subject factor and MDL subregion as a within-subject factor revealed a significant interaction between cortical target and MDL position (*F*(2, 26) = 25.84, *p* < 0.000). Tukey adjusted post hoc comparisons show that PRL-projecting neurons were concentrated in anterior MDL (Anterior: 334 ± 42.3 cells/mm^2^; Central: 179 ± 36.6; Posterior: 119 ± 36.9; anterior > central: *p* = 0.001; anterior > posterior: *p* < 0.000), whereas ACC-projecting neurons were enriched in posterior MDL (Posterior: 280 ± 58.1 cells/mm^2^; Central: 176 ± 33.5; Anterior: 112 ± 25.6; posterior > anterior: *p* = 0.000; posterior > central: *p* = 0.022). (D) Retrograde tracer (CTb) injections into anterior MDL (n = 4) or posterior MDL (n = 4). Representative tissue images show labeled neurons in (left to right) PRL and ACC. (E) Nissl atlas images from the Allen Reference Atlas - Mouse Brain showing PRL and ACC cell quantification boundaries. (F) Anterior MDL injections preferentially labeled PRL neurons (PRL: 154.0 ± 14.1 cells/mm^2^; ACC: 18.7 ± 14.1; *t*(6) = 7.19, *p* < 0.001), while posterior MDL injections preferentially labeled ACC neurons (ACC: 149.5 ± 14.1 cells/mm^2^; PRL: 8.6 ± 14.1; *t*(6) = −7.49, *p* < 0.001). Linear mixed-effects ANOVA confirmed a significant MDL injection × PFC region interaction (*F*(1, 6) = 107.75, *p* < 0.001), with no significant main effects of injection site (*F*(1, 6) = 0.24, *p* = 0.640) or cortical target (*F*(1, 6) = 0.04, *p* = 0.840).

The anterior-posterior segregation of MD subpopulations and their corresponding cortical inputs suggest that the MD could support region-specific computations within the PFC through recruitment of specific output pathways. While previous lesion and inactivation studies demonstrate that the MD is required for predictive cues to guide behavior (Corbit & Balleine, 2003; Fisher et al., 2020; Parnaudeau et al., 2015), these studies have not assessed the possibility that the anatomically dissociable circuits identified above are differentially engaged in this process. Therefore, we measured changes in calcium activity in MD terminals in the ACC or PRL using fiber photometry as animals underwent Pavlovian cue-reward pairing. MD→ACC and MD→PRL calcium activity at key timepoints: early training when the cue is novel (Day 1) and late training after the cue has been associated with reward (Day 11; Figure 3A, F). Analyses of MD axonal Ca^+2^ activity focused on (1) change in signal amplitude in response to cue presentation (defined as the difference between the peak signal (in peak ΔF/F) following cue onset and the pre-cue baseline, (2) area under the curve (AUC) of the Ca^+2^ timeseries during the 5s following cue onset, and (3) the time course of Ca^+2^ changes, defined as the time in seconds required for the cue-evoked peak to decay to 10% of its maximum amplitude (t90).

**Figure 3.**
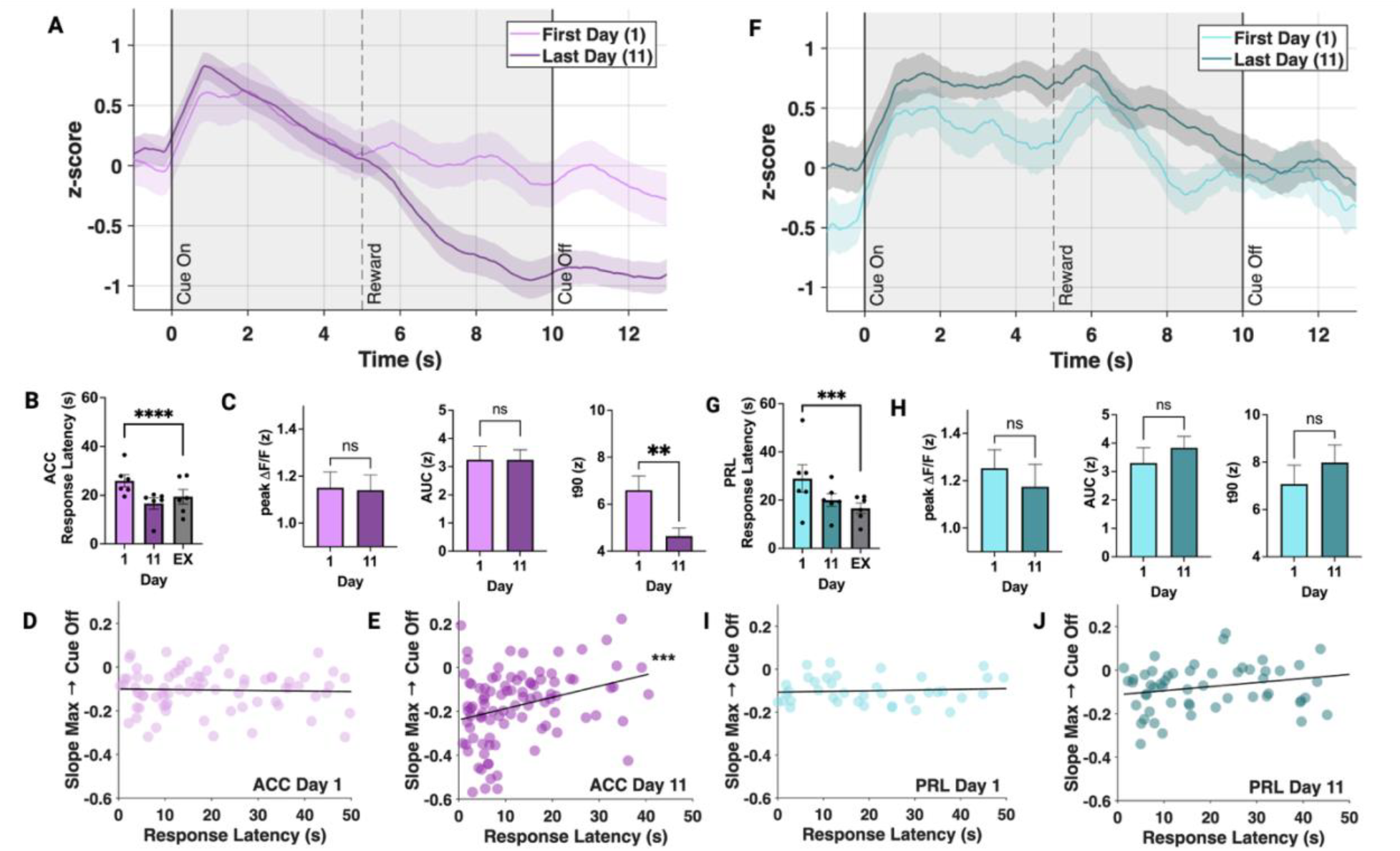
MD→PRL and MD→ACC terminal activity differentiates early vs late learning and tracks cue-guided responding. (A) MD terminals in ACC average trace Day 1 and Day 11 (B) Average response latency on Day 1, 11, Extinction (Day 1 vs. Extinction, *t*(372) = 5.592, *p* < 0.000) (C) ACC group signal features, Day 1 vs. Day 11; peak ΔF/F (Day 1 mean = 1.17 ± 0.142; Day 11 mean = 1.21 ± 0.137; *t*(413) = -0.433, *p* = 0.902); AUC (Day 1 mean = 3.05 ± 0.569; Day 11 mean = 3.34 ± 0.554; *t*(420) = -0.550, *p* = 0.847); t90 (Day 1 mean = 6.59 ± 0.650; Day 11 mean = 4.60 ± 0.591; *t*(291) = 2.940, *p* = 0.010). (D) ACC Day 1 Slope vs latency (*r* = -0.05, *p* = 0.642) (E) ACC Day 11 Slope vs latency (*r* = 0.31, *p* = 0.002) (F) MD terminals in PRL average trace Day 1 and Day 11 (G) Average response latency on Day 1, 11, Extinction (Tukey corrected post-hoc tests; Day 1 vs. Extinction, *t*(371) = 3.714, *p* = 0.001) (H) PRL group signal features, Day 1 vs. Day 11; peak ΔF/F (Day 1 mean = 1.50 ± 0.163; Day 11 mean = 1.35 ± 0.154; *t*(411) = 1.068, *p* = 0.534); AUC (Day 1 mean = 3.55 ± 0.706; Day 11 mean = 4.01 ± 0.650; *t*(420) = -0.715, *p* = 0.755); t90 (Day 1 mean = 7.12 ± 0.787; Day 11 mean = 8.54 ± 0.751; *t*(287) = -1.619, *p* = 0.239). (I) PRL Day 1 Slope vs latency (*r* = 0.12, *p* = 0.455) (J) PRL Day 11 Slope vs latency (*r* = 0.21, *p* = 0.151)

We first examined learning-related changes in MD→ACC terminal activity. Across all days of training, presentation of the reward predictive cue (overhead light) evoked an increase in axonal Ca^2+^ concentrations. The magnitude of this cue-evoked increase was consistent across Days 1 and 11 (Figure 3C). However, calcium signals in MD→ACC terminals showed a markedly different temporal profile across training days, characterized by stronger suppression and faster decay following the peak on Day 1 vs Day 11 (Figure 3C). Although MD→ACC amplitude-related measures (peak ΔF/F and AUC) showed relatively modest changes across days, signal decay dynamics were strongly affected by learning: t90 exhibited a robust effect of Day (*F*(2, 271.32) = 9.69, *p* = 8.59 × 10^−5^). This indicates that, in ACC, learning primarily reshapes the timing of MD terminal responses rather than their overall magnitude. To address this possibility explicitly, we quantified the relationship between trial-by-trial slopes (calculated from the maximum post-cue onset to the end of the CS) and animals’ response latency. We found that Ca^2+^ decay rates correlated with response latencies on Day 11 (Figure 3D), but not on Day 1 (Figure 3E). Together, our data suggests that suppression of cue-evoked MD→ACC activity reflects the decision to use the cue to guide behavior after learning the predictive relationship between the two. These results are also in line with previous evidence that the MD plays a role in controlling the timing of actions (Lusk et al., 2020).

In contrast to the pathway to the ACC, MD→PRL terminal activity showed little modification as a function of training. On both days of interest (Days 1 and 11; Figure 3F), calcium activity in MD terminals in PRL increased following cue onset, remained elevated until reward delivery, and peaked near reward delivery. Neither cue evoked amplitude, AUC, or the overall time course of axonal Ca^2+^ concentrations differed significantly between Days 1 and 11 (Figure 3H), indicating that the PRL response was a stable phenotype largely independent of learning-related changes. Moreover, PRL terminal slopes were not correlated with response latency on either day (Day 1: Figure 3I; Day 11: Figure 3J), further suggesting that MD→PRL activity does not track behavioral performance in the same way as MD→ACC activity. Importantly, average response latency decreased significantly across testing for both groups (Figure 3B, 3G), indicating similar overall improvements in performance and task engagement. Thus, these regional differences in MD projection recruitment reflect stable, circuit-specific signatures of recruitment during cue-guided behavior.

These results indicate that learning preferentially reshapes the temporal dynamics of MD→ACC terminal responses, while MD→PRL responses remain comparatively stable. Thus, MD→ACC projections develop a selective, learning-dependent, and behavior-predictive role in guiding cue-driven behavior, whereas MD→PRL projections exhibit more modest or transient modulation.

Finally, we examined MD terminal responses during an extinction session in which the predictive cue was no longer followed by pellet delivery (Figure 4), providing a window into the regulation of these circuits when outcomes violate previously learned associations (Alcaraz et al., 2018). MD→ACC terminals displayed a markedly different pattern during extinction relative to task training days (Figure 4A). There was a significant increase in MD→ACC recruitment as measured by peak and AUC during late extinction trials vs early extinction trials and Day 11 (peak ΔF/F (Early: 1.10 ± 0.186; Late: 1.49 ± 0.163; *t*(120) = 2.350, *p* = 0.020); AUC (Early: 2.86 ± 0.915; Late: 4.51 ± 0.837; *t*(119) = 2.367, *p* = 0.019); Figure 4B). Further, the timecourse of cue-evoked activity, as measured by t90, was consistent across extinction training (Figure 4B), however interestingly, the relationship between the slope of the decline of axonal Ca^2+^ and reward approach latency was lost (Figure 4C). There was no difference in response behavior between trial groups (Figure 4D). These patterns of responsivity across training extinction suggest that neural circuitry associated with encoding the outcome of cue-based predictions may have an inhibitory influence on the MD-ACC pathway. In the absence of such feedback-related inhibition, prolonged activation of the MD-ACC circuit encodes a violation of expectation (Brown & Braver, 2005; Cole et al., 2024).

**Figure 4.**
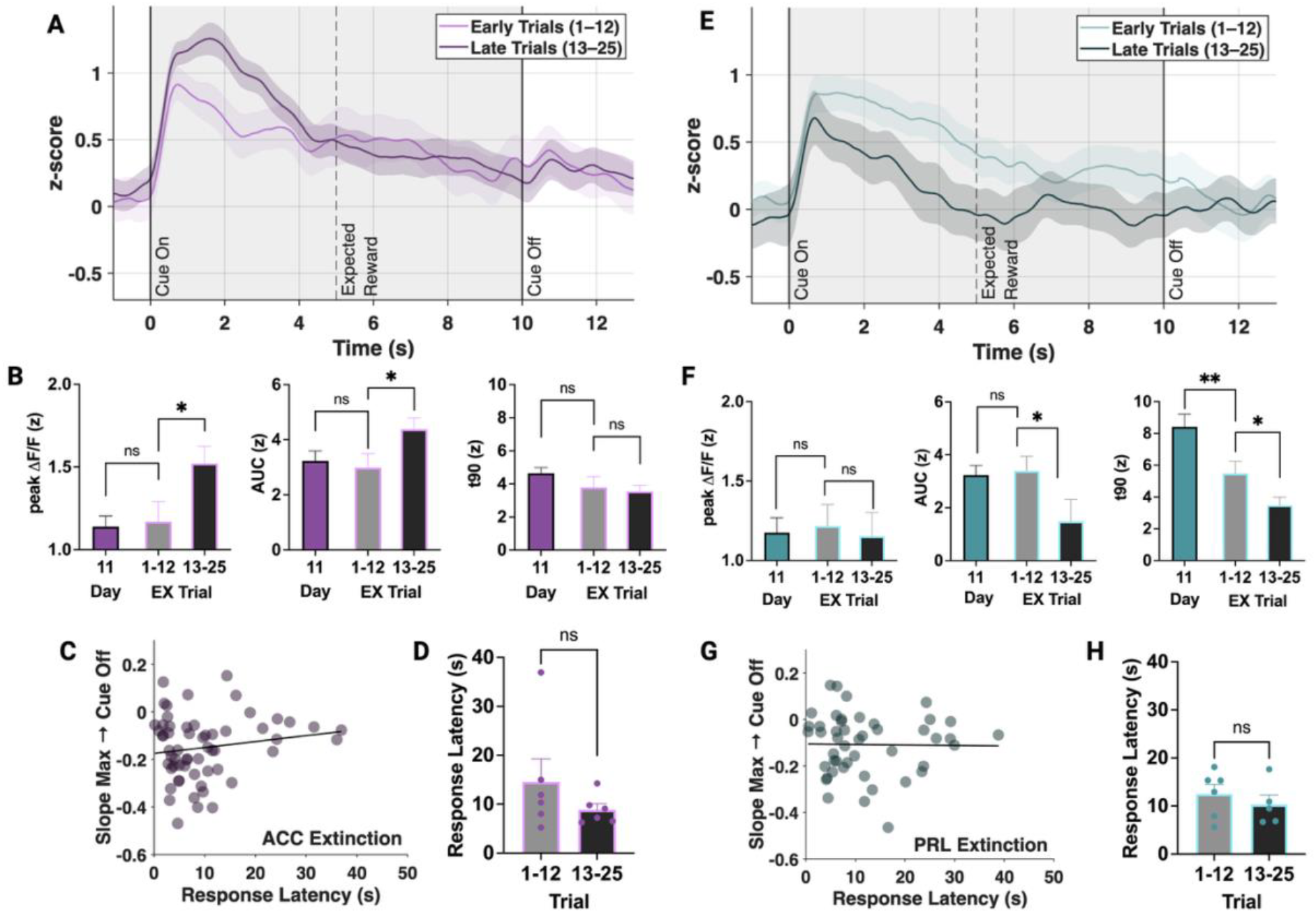
MD terminals in PRL and ACC show opposing activity during cue-reward extinction. (A) ACC average trace of Extinction trial groups (1-12, 13-25) (B) ACC group signal features, Day 11 vs. Early extinction trials; peak ΔF/F (Day 11: 1.23 ± 0.143; *t*(206) = -0.369, *p* = 0.712); AUC (Day 11: 3.31 ± 0.420; *t*(204) = -0.320, *p* = 0.749); t90 (Day 11: 4.63 ± 0.503; *t*(148) = -0.806, *p* = 0.421). Extinction Day trials 1-12 vs. trials 13-25; peak ΔF/F (Early: 1.10 ± 0.186; Late: 1.49 ± 0.163; *t*(120) = 2.350, *p* = 0.020); AUC (Early: 2.86 ± 0.915; Late: 4.51 ± 0.837; *t*(119) = 2.367, *p* = 0.019); t90 (Early: 3.90 ± 0.919; Late: 3.73 ± 0.774; *t*(82.7) = -0.216, *p* = 0.830). (C) ACC Extinction Day Slope vs. response latency (*r* = 0.16, *p* = 0.219) (D) ACC Response latency early vs. late trials (*t*(67.7) = -0.726, *p* = 0.470) (E) PRL average trace of Extinction trial groups (1-12, 13-25) (F) PRL group signal features, Day 11 vs. Early extinction trials; peak ΔF/F (Day 11: 1.38 ± 0.163; *t*(199) = -1.271, *p* = 0.205); AUC (Day 11: 3.93 ± 0.522; *t*(162) = -0.782, *p* = 0.435); t90 (Day 11: 8.33 ± 0.672; *t*(86.1) = -2.869, *p* = 0.005). Extinction Day trials 1-12 vs. trials 13-25; peak ΔF/F (Early: 1.17 ± 0.188; Late: 1.18 ± 0.192; *t*(120) = 0.048, *p* = 0.961); AUC (Early: 3.82 ± 0.933; Late: 2.03 ± 0.954; *t*(119) = -2.367, *p* = 0.019); t90 (Early: 5.71 ± 0.889; Late: 4.86 ± 0.940; *t*(85.3) = -2.049, *p* = 0.044). (G) PRL Extinction Day Slope vs. response latency (*r* = -0.01, *p* = 0.923) (H) PRL Response latency early vs. late trials (*t*(69.0) = -0.162, *p* = 0.872) (I) ACC average trace of Extinction trial groups (1-12, 13-25) (J) ACC group signal features, Day 11 vs. Early extinction trials; peak ΔF/F (Day 11: 1.23 ± 0.143; *t*(206) = -0.369, *p* = 0.712); AUC (Day 11: 3.31 ± 0.420; *t*(204) = -0.320, *p* = 0.749); t90 (Day 11: 4.63 ± 0.503; *t*(148) = -0.806, *p* = 0.421). Extinction Day trials 1-12 vs. trials 13-25; peak ΔF/F (Early: 1.10 ± 0.186; Late: 1.49 ± 0.163; *t*(120) = 2.350, *p* = 0.020); AUC (Early: 2.86 ± 0.915; Late: 4.51 ± 0.837; *t*(119) = 2.367, *p* = 0.019); t90 (Early: 3.90 ± 0.919; Late: 3.73 ± 0.774; *t*(82.7) = -0.216, *p* = 0.830). (K) ACC Extinction Day Slope vs. response latency (*r* = 0.16, *p* = 0.219) (L) ACC Response latency early vs. late trials (*t*(67.7) = -0.726, *p* = 0.470)

In contrast, there was no significant difference in cue-evoked signal peak between Day 11 and early (trials 1-12) or late (13-25) trials during the extinction session in PRL. However, we noted that the AUC of cue evoked activity decreased significantly by the last half of the extinction session (Figure 4E, F). These data suggest that although the cue still triggered an initial response in the MD-PRL pathway, this activation became weaker as the cue lost its predictive validity. Consistent with this, cue-evoked MD→PRL activity no longer remained elevated until the expected time of reward delivery, but instead showed a rapid suppression, as reflected by a decrease in t90. Importantly, there was no difference in response behavior between trial groups (Figure 4H) and the relationship between the slope of the decline of MD→PRL activity did not correlate with latency during the extinction session (Figure 4G), suggesting that these changes in overall recruitment of the MD-PRL pathway did not reflect more general changes in extinction-related learning.

## Discussion

The present anatomical findings reveal a striking anterior–posterior organization within the mediodorsal thalamus that maps onto distinct subdivisions of prefrontal cortex. MD→PRL projections arise primarily from anterior MDL, whereas MD→ACC projections originate from posterior MDL. Complementary retrograde tracing from cortex back into MD demonstrates the same spatial specificity, indicating that MD-PFC communication is organized into reciprocal, segregated loops rather than a single convergent pathway.

This architecture has important implications for theories of thalamocortical function. Contemporary models describe the MD as a higher-order thalamic nucleus that supports PFC-dependent cognition by coordinating cortical activity during learning and attentional control (Mitchell, 2015; Schmitt et al., 2017). Our data refine this framework by demonstrating that MD output is not uniform across PFC. Instead, distinct MD subcircuits interface with functionally specialized cortical territories: MD projections to PRL are uniquely engaged by cues that serve as reliable predictors of appetitive outcomes, receives input from anterior MDL. MD projections to the ACC are implicated in outcome integration and receive input from posterior MDL. The presence of reciprocal loops suggests that these interactions support dynamic, bidirectional modulation of network states during behavior.

Importantly, this circuit organization has translational implications. Disruption of MD-PFC connectivity is a hallmark feature of neuropsychiatric conditions characterized by deficits in executive function, including schizophrenia, ADHD, and depression (Anticevic et al., 2014; Minzenberg et al., 2009). The fact that MD-PFC interactions are anatomically segregated suggests that disease-related impairments may arise from selective vulnerability of specific loops rather than global thalamocortical dysfunction. Consistent with this anatomical segregation, our functional data reveal pathway-specific dynamics during learning and extinction. These findings suggest that cognitive symptoms may reflect dysfunction in specific thalamocortical loops rather than a unitary deficit in prefrontal control. For example, alterations in posterior MDL-ACC connectivity could contribute to impaired error monitoring or difficulty evaluating feedback, whereas disruptions in anterior MDL-PRL loops may affect selective attention. Understanding the precise organization of these pathways is therefore essential for identifying more targeted mechanisms underlying symptoms.

One potential mechanism underlying these divergent response profiles is target-specific inhibition. ACC-projecting MD neurons may be selectively modulated by inhibitory circuits, such as the thalamic reticular nucleus or local cortical interneurons, which could suppress or shape their activity in a context-dependent manner. In contrast, PRL-projecting MD neurons may experience less inhibitory modulation, allowing more stable signaling across task phases. This selective inhibition could explain why MD-ACC activity flexibly tracks cue reliability, while MD-PRL signals remain stable but fall during cue-reward extinction and suggests that circuit-specific disruptions could produce distinct cognitive deficits depending on which thalamocortical loop is affected. Understanding these mechanisms may help guide targeted interventions aimed at restoring the function of discrete thalamocortical loops in neuropsychiatric disorders.

More broadly, these findings highlight how the brain can organize information flow in a highly specific and modular way. Rather than sending the same signal everywhere, the MD output neurons are tailored to support the functions of their cortical target. This modular architecture may help the brain handle complex behaviors that require attending to multiple cues, adjusting strategies on the fly, or learning new rules without interfering with ongoing task performance. It also emphasizes that attention, learning, and decision-making are not global brain states, but the result of many finely tuned circuits interacting in parallel. By uncovering this structural logic, our work provides a framework for understanding how discrete thalamocortical loops can support flexible, goal-directed behavior in both healthy and diseased brains.

## Materials and Methods

### Mice

Adult male and female C57BL/6J mice (>90 days old) housed in standard laboratory cages were used in the present experiments. Separate cohorts of mice were used for fiber photometry (n = 12) and circuit tracing experiments (n = 23). All procedures were conducted in adherence with protocols approved by the Institutional Animal Care and Use Committee of Virginia Tech. For fiber photometry experiments, animals were single-housed and trained daily over an approximately 21-day experimental period. Water was available *ad libitum* and food (regular chow) was measured out daily (approximately 3g per day for males, 2.5g per day for females) to maintain 90% of their individual free-feeding weight. Animals were kept on a reverse light/dark cycle, and all training and testing took place during the dark cycle between 9:00 A.M. and 9:00 P.M.

### Surgery: Fiber photometry experiments

Prior to shaping and training on the task, all mice underwent stereotaxic surgeries. Mice were anesthetized with isoflurane (1-4%) and then positioned on a stereotaxic apparatus (RWD Life Science, China). An injection of jGcAMP7s (pGP-AAV-syn-jGCaMP7s-WPRE; Addgene; titer ≥ 1 × 10^13^ vg/mL) was administered into the mediodorsal thalamus (MD; AP -1.30 mm; ML +/-0.45 mm; DV -3.3 mm) at a volume of 700 nL and a rate of 100nl/min using a Hamilton syringe. The injection needle was left in place for 5 min to allow diffusion. In the same surgery, a 400µm optical fiber (Doric Lenses) was implanted unilaterally into either the prelimbic cortex (PRL; AP +1.78 mm; ML +/-0.3 mm; DV -2.3 mm) or anterior cingulate cortex (ACC; AP +1.18 mm; ML +/-0.4 mm; DV -1.7 mm). Animals were monitored closely for 5 days post-surgery and were allowed to recover for a minimum of 3 weeks before behavioral experiments. Fiber placements and viral expression were verified histologically, and animals with incorrect placements were excluded.

### Fiber Photometry recording and analysis

Fiber photometry was used to record calcium activity from mediodorsal thalamus (MD) axon terminals in the prefrontal cortex of freely moving mice. Following post-surgical recovery, animals were handled for 20 min per day for five consecutive days and subsequently habituated to the fiber optic patch cable in the behavioral chamber for three days (30 min/day). Recordings were performed in a Noldus behavioral chamber equipped with a house light and reward port using a fiber photometry system (Doric Lenses) and Synapse software (Tucker-Davis Technologies). Excitation light at 488 nm was used to measure calcium-dependent fluorescence, while an isosbestic control signal (405 nm), which produces calcium-independent fluorescence, was recorded simultaneously to control for motion artifacts, photobleaching, and other non-calcium-dependent changes in fluorescence. Photometry data were processed in MATLAB by fitting the isosbestic control signal to the calcium-dependent channel using linear regression and calculating the normalized fluorescence change (ΔF/F; ΔF/F = (F488 − F405-fit) /F405-fit). Signals were acquired at 1017 Hz, downsampled to match the EthoVision video acquisition rate (29.97 Hz), and smoothed using a 500 ms moving window. Signals were demodulated in Synapse prior to analysis. To enable comparisons across animals and sessions, fluorescence signals within each training session were normalized using z-score transformation. Statistical analyses examining effects of region and training day were conducted using linear mixed-effects models in R Studio.

### Behavioral recordings and analysis

The experiments were designed to measure cue-evoked behavioral responses during a visual cue detection task. To do this, population-level calcium dynamics of MD axon terminal fields were recorded in the prefrontal cortex (PRL or ACC). To measure cue detection and utilization, a classical visual cue detection task was used in these experiments. Animals underwent approximately one week of habituation to the testing chamber (Noldus) and patch cord, as well as food shaping to learn where the reward will be dispensed from. Once the animal ate over 50% of the available sugar pellets for two days in a row, training in the cue detection task began. Each training day consisted of 25 trials with an inter-trial interval of 90 ± 30 s. A trial begins with the illumination of the chamber’s house light which remains on for 10s. In the middle of this cue light (at 5s), a sugar pellet reward is dispensed into the reward port. An infrared beam inside the reward port monitored nose pokes. The primary behavioral variable was latency to approach the reward port, expressed in seconds from cue onset. This task does not explicitly require the use of the cue to receive the reward. Instead, reward delivery was independent of the animal’s response (measured by a nose poke into the reward port). Trials in which animals did not enter the reward port before the end of the inter-trial interval were classified as omissions and excluded from latency and photometry analyses. During extinction sessions, the cue light was presented without reward delivery.

### Surgery and quantification: Circuit tracing experiments

Adult male and female C57BL/6J mice above 90 days of age were used in the present experiments (n = 23). For experiment 1, 100 nL of Cholera Toxin Subunit B (CTb; Recombinant), Alexa Fluor 647 Conjugate (Thermo Fisher) was injected into either the PRL (AP +1.78 mm; ML +/-0.3 mm; DV -2.3 mm) or the ACC (AP +1.18 mm; ML +/-0.4 mm; DV -1.7 mm). For experiment 2 tracing PFC to MD pathways, 100 nL of CTb was injected into either the anterior MDL (AP -1.06 mm; ML +/-0.7 mm; DV -3.3 mm) or posterior MDL (AP -1.70 mm; ML +/-0.7 mm; DV -3.2 mm). Animals were transcardially perfused with PBS followed by 4% paraformaldehyde 7 days after injection. Brains were sliced at 40 µm using a microtome, and tissue was imaged using a Nikon Confocal microscope (60X magnification). Cell density (cells/mm^2^) was calculated by dividing the number of labeled cells by the area of a standardized ROI defined for each region using ImageJ/Fiji.

### Statistical analysis

Cell count data were analyzed using linear mixed-effects models and repeated-measures ANOVA in R. Two complementary analyses were performed. First, to assess the distribution of MDL cell bodies as a function of cortical injection site, cell counts across anterior-posterior MDL subregions were analyzed with injection site (PRL vs ACC) as a between-subject factor and MDL subregion as a within-subject factor. Subject was included as a random effect to account for repeated measurements within animals. Repeated-measures ANOVA was performed using mixed-effects modeling with Kenward-Roger corrected degrees of freedom, followed by Tukey-adjusted post hoc comparisons of estimated marginal means. Second, to examine cortical cell body counts as a function of MDL injection location, cell counts in PRL and ACC were analyzed with MDL injection site (anterior vs posterior) as a between-subject factor and cortical region (PRL vs ACC) as a within-subject factor. Linear mixed-effects models were fit with Subject included as a random intercept. Main effects and interactions were assessed using ANOVA, and follow-up comparisons were conducted using estimated marginal means with Tukey adjustment.

Each feature extracted from the photometry data was analyzed using linear mixed-effects models with experimental Day and brain region (PRL, ACC) included as fixed effects and Animal ID included as a random intercept to account for repeated measurements within subjects. For models including both regions, Day × Region interactions were tested. Statistical significance of fixed effects was assessed using Type III ANOVA with Kenward-Roger degrees-of-freedom correction. Region-specific effects of Day were further examined using separate LMEMs for PRL and ACC, followed by post hoc pairwise comparisons of estimated marginal means with Tukey correction for multiple comparisons.

